# Toxin-producing endosymbionts shield pathogenic fungus against micropredators

**DOI:** 10.1101/2022.01.07.475374

**Authors:** Ingrid Richter, Silvia Radosa, Zoltán Cseresnyés, Iuliia Ferling, Hannah Büttner, Sarah P. Niehs, Ruman Gerst, Marc Thilo Figge, Falk Hillmann, Christian Hertweck

## Abstract

The phytopathogenic fungus *Rhizopus microsporus* harbours a bacterial endosymbiont (*Mycetohabitans rhizoxinica*) for the production of the toxin rhizoxin, the causative agent of rice seedling blight. This toxinogenic bacterial-fungal alliance is, however, not restricted to the plant disease, but has been detected in numerous environmental isolates from geographically distinct sites covering all five continents. Yet, the ecological role of rhizoxin beyond rice seedling blight has been unknown.

Here we show that rhizoxin serves the fungal host in fending off protozoan and metazoan predators. Fluorescence microscopy and co-culture experiments with the fungivorous amoeba *Protostelium aurantium* revealed that ingestion of *R. microsporus* spores is toxic to *P. aurantium*. This amoebicidal effect is caused by the bacterial rhizoxin congener rhizoxin S2, which is also lethal towards the model nematode *Caenorhabditis elegans*. By combining stereomicroscopy, automated image analyses, and quantification of nematode movement we show that the fungivorous nematode *Aphelenchus avenae* actively feeds on *R. microsporus* that is lacking endosymbionts, while worms co-incubated with symbiotic *R. microsporus* are significantly less lively.

This work uncovers an unexpected ecological role of rhizoxin as shield against micropredators. This finding suggests that predators may function an evolutionary driving force to maintain toxin-producing endosymbionts in non-pathogenic fungi.

## 1 Introduction

The filamentous fungus *Rhizopus microsporus* (Phylum: Mucoromycota) plays an important role in a variety of fields including agriculture, biotechnology, and medicine. While some strains are being used for food fermentation and metabolite production others cause mucormycosis in immunocompromised patients (Lackner, Partida-Martinez, et al., 2009; Scherlach et al., 2013). However, *R. microsporus* has gained the most attention as the causative agent of rice seedling blight, a plant disease that causes severe crop losses in agriculture in Asia (Lackner & Hertweck, 2011). The disease is mediated through the highly potent phytotoxin rhizoxin (Figure 1), which efficiently stalls plant cell division by binding to the β-tubulin of the rice plant cells. This leads to abnormal swelling of the tips of rice seedling roots, eventually leading to plant death (Iwasaki et al., 1984). While *R. microsporus* was initially believed to be the toxin producer, we discovered that rhizoxin is biosynthesised by endosymbiotic beta-Proteobacteria, *Mycetohabitans rhizoxinica*, residing within the fungal hyphae (Partida-Martinez & Hertweck, 2005). *M. rhizoxinica* does not only provide the fungus with potent toxins, but also regulates fungal reproduction (Mondo et al., 2017; Partida-Martinez, Monajembashi, et al., 2007). This toxinogenic bacterial-fungal alliance is globally distributed across all five continents inhabiting a variety of niches ranging from temperate and arid soils to human tissue (Kerr et al., 1988; Lackner, Mobius, et al., 2009). In one of these eight toxin-producing *Rhizopus-Mycetohabitans* strains rhizoxin was shown to be a potent phytotoxin (Figure 1), while the ecological role of rhizoxin in the other *Rhizopus* strains is currently unknown.

**Figure 1.**
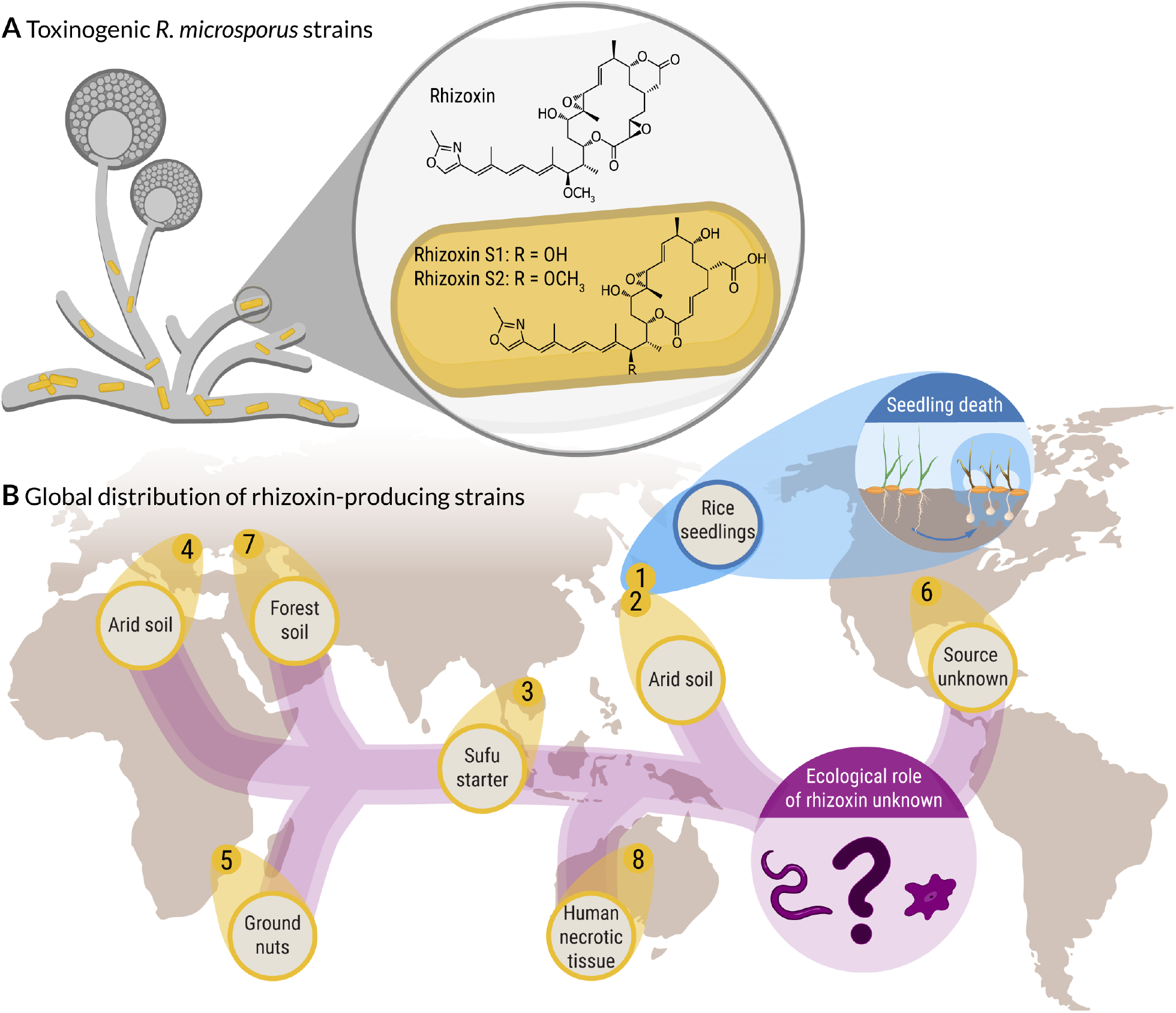
Global distribution of a toxin-producing bacterial-fungal symbiosis. **(A)** Symbiotic bacteria (*Mycetohabitans* sp.), residing within the fungal hypha of *Rhizopus microsporus*, produce a mixture of toxic secondary metabolites (rhizoxins). **(B)** Rhizoxin-producing *Rhizopus-Mycetohabitans* strains were isolated from environmental samples from geographically distinct sites covering all five continents. In one of the eight toxinogenic strains (*Rhizopus microsporus* ATCC 62417, blue), rhizoxin causes blight disease in rice seedlings, while the ecological role of rhizoxin in the other, non-pathogenic *Rhizopus* strains is currently unknown.

Since fungi are able to utilise toxic secondary metabolites to protect themselves from predators and antagonistic organisms (Boysen et al., 2021; Künzler, 2018; Spiteller, 2015), we reason that rhizoxin might act as an anti-predator agent in non-pathogenic *Rhizopus* strains. Effective defence strategies are particularly important for fungi as their high nutrient content, large biomass, and inability to move, makes fungi an ideal food source for micropredators such as soil-dwelling amoeba, nematodes, mites, and springtails (Baumann, 2018; Fierer, 2017; Fountain & Hopkin, 2005; Radosa et al., 2019; Radosa et al., 2021; Ruess & Lussenhop, 2005; Yeates et al., 1993). For example, the soil mould *Aspergillus nidulans* relies on secondary metabolites to defend itself against the fungivorous springtail *Folsomia candida* (Döll et al., 2013; Rohlfs et al., 2007), while aflatoxin protects *Aspergillus flavus* from fungivory by insects (Drott et al., 2017).

Whilst most studies focus on species belonging to the Ascomycota and Basidiomycota, reports on toxic defence molecules produced by Mucoromycota fungi are scarce. Interestingly, Mucoromycota fungi often harbour endobacteria (Bonfante & Venice, 2020; Okrasińska et al., 2021), which can produce toxic secondary metabolites that shield the fungal host from predatory nematodes (Büttner et al., 2021). This strategy to fend off predators might be a common trait in symbiotic Mucoromycota fungi, as all toxic compounds identified in Mucoromycota fungi so far are produced by endofungal bacteria. For example, *M. rhizoxinica* is a fungal endosymbiont with a remarkable potential to produce secondary metabolites (Niehs et al., 2020; Niehs et al., 2019; Niehs, Dose, et al., 2018; Niehs, Scherlach, et al., 2018), despite its small genome size (3.75 Mb) (Lackner et al., 2011). Among the many secondary metabolites produced by *M. rhizoxinica*, rhizoxin represents a prime candidate as a potential anti-predator agent. Rhizoxin exhibits its effect against most eukaryotes including vertebrates and fungi by efficiently binding to β-tubulin, which causes disruption of microtubule formation (Schmitt et al., 2008; Takahashi et al., 1987).

However, apart from being a potent phytotoxin, the ecological role of rhizoxin is still unclear and raises the question why soil-borne fungi harbour toxin-producing bacteria. Here, we tested the effects of rhizoxin on the model nematode *Caenorhabditis elegans*, and two mycophagous eukaryotes, the amoeba *Protostelium aurantium* and the nematode *Aphelenchus avenae*. We show that rhizoxin-producing endosymbionts can prevent killing of *R. microsporus* by protozoan and metazoan fungivorous micropredators.

## 2 Results

### 2.1 Ingestion of *R. microsporus* spores is toxic to a fungivorous amoeba

Within the soil community, fungi are constantly challenged by antagonistic organisms such as amoebae, which are well known for their micropredatory lifestyle (Novohradská et al., 2017). Using the recently isolated amoeba *Protostelium aurantium*, an amoeba that specifically feeds on fungi either by phagocytosis of yeastlike cells or by invasion of mature hyphae (Hillmann et al., 2018; Radosa et al., 2019), we investigated if *R. microsporus* endosymbionts are able to protect their fungal host from this predatory amoeba.

In a co-culture experiment, *P. aurantium* was incubated with either dormant or swollen spores of *R. microsporus* (ATCC62417). Dormant spores are readily ingested by *P. aurantium*, while swollen spores are taken up less frequently (Figure 2A—video supplement 1). This difference can be explained by the size of the swollen spores (9.1 μm ± 0.9 μm, Figure supplement 1), which are significantly larger than dormant spores (5.2 μm ± 0.5 μm, unpaired *t*-test: *t* = 5.93, df = 3.2, *p* = *0.0081*—Supplementary file 1). A reduced uptake of swollen *R. microsporus* spores was previously also reported for macrophages (Itabangi et al., 2020). Although the increase in spore size makes it difficult for *P. aurantium* (mean diameter of approximately 13.2 μm ± 2.2 μm) to ingest swollen spores, in some rare cases we observed phagocytosis of swollen spores and *R. microsporus* germlings (Figure supplement 2). Survival assays revealed that the presence of spores reduces the viability of the amoebae, with dormant spores being significantly more harmful for *P. aurantium* than swollen spores (unpaired *t*-test: *t* = 6.33, df = 2.6, *p* = *0.0122*, Figure 2B—Supplementary file 2). This sensitivity correlates with a higher frequency of phagocytosis for dormant spores of *R. microsporus*.

**Figure 2.**
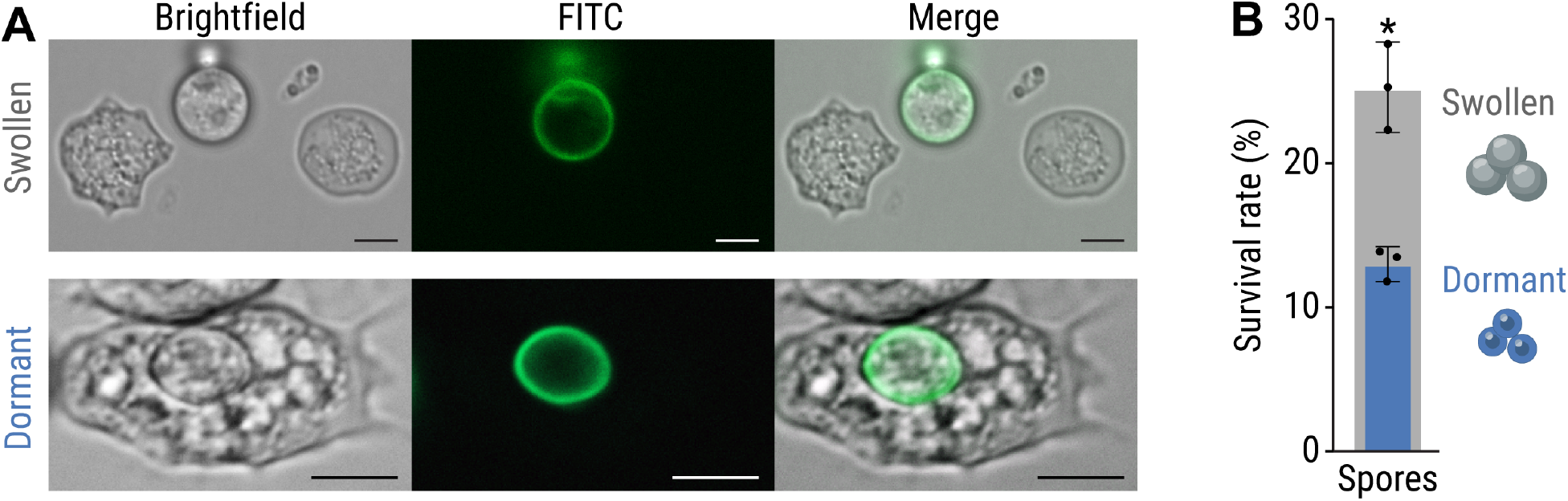
Predation of *Protostelium aurantium* on spores of *Rhizopus microsporus*. **(A)** Fluorescence microscopy images showing FITC-stained, swollen *R. microsporus* spores (top) and ingestion of a dormant *R. microsporus* spore by *P. aurantium* (bottom). Scale bars: 5 μm. **(B)** Feeding of *P. aurantium* on dormant spores (blue) leads to a reduced survival rate of *P. aurantium* compared to swollen spores (grey). N = 3 independent replicated experiments ± one SEM. Unpaired *t*-test with Welch’s correction (\**p<0.05*, Supplementary file 2).

### 2.2 Endosymbionts protect fungal host from amoeba predation

To test whether bacterial endosymbionts are responsible for amoeba killing, trophozoites of *P. aurantium* were exposed to the following fungal culture extracts: (i) symbiotic (endosymbiont-containing) *R. microsporus* ATCC62417 (RMsym); and (ii) apo-symbiotic *R. microsporus* ATCC62417/S (RMapo). Amoebae were co-incubated with either 2% or 5% of the culture extracts for 1 hr. Following exposure, the treated samples were placed on agar plates containing *Rhodotorula mucilaginosa* as a food source. Living amoebae, incubated with solvent control, form a visible predation plaque (clearance of yeast) that expands over five days of incubation (Figure 3A,B). Increasing predation plaques also appeared when amoeba were incubated with extracts from apo-symbiotic *R. microsporus* hyphae but predation plaques were not detected when incubated with crude extracts from symbiotic *R. microsporus (p<0.0001*, Figure 3A,B—Supplementary file 3). This confirms that *P. aurantium* is inhibited or killed due to the presence of metabolites produced by endosymbionts.

**Figure 3.**
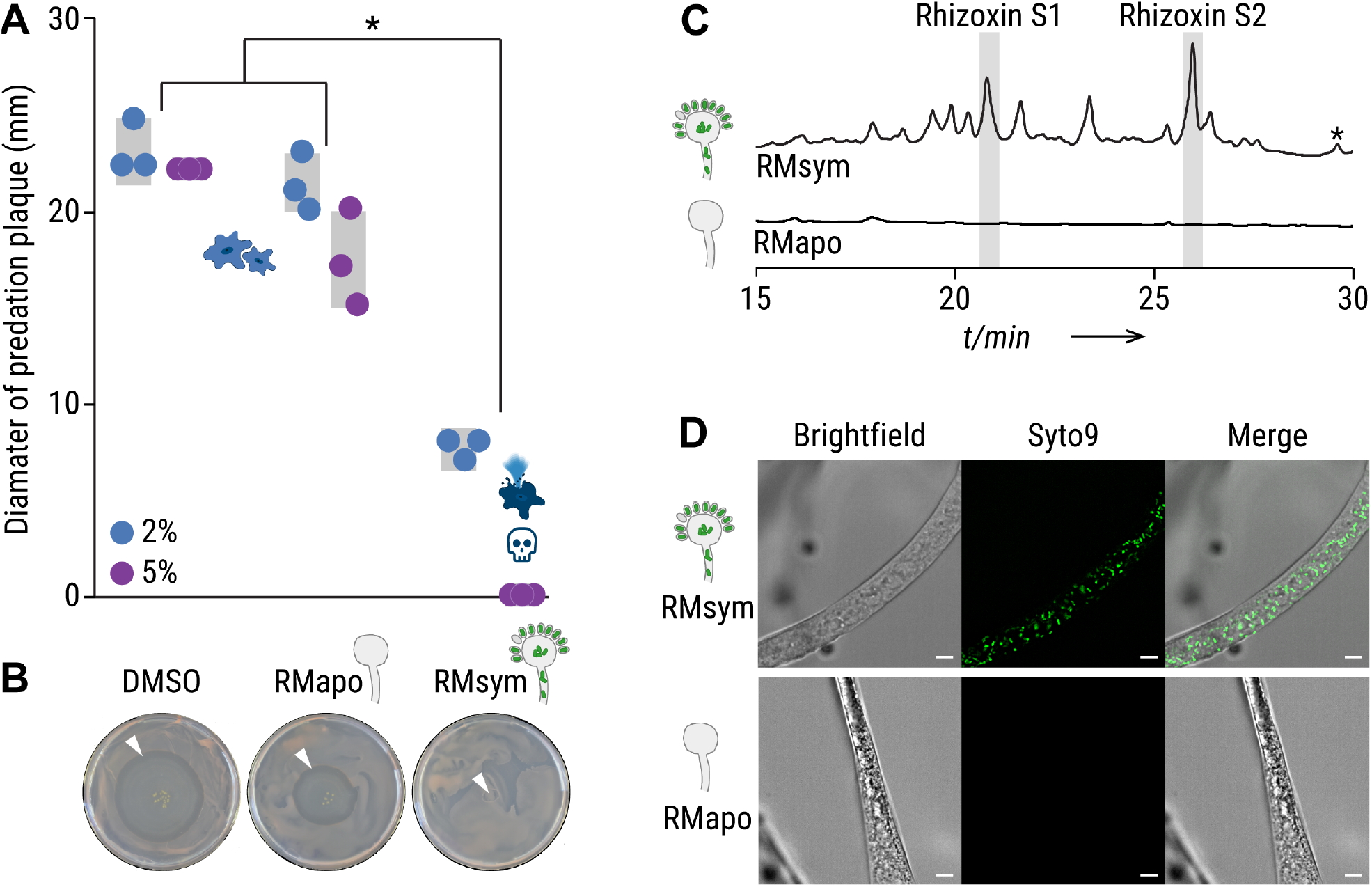
Culture extracts from symbiotic *Rhizopus microsporus* kills *Protostelium aurantium*. **(A)** The viability of *P. aurantium*, indicated by the diameter of the predation plaque (clearance of yeast), is significantly reduced in cultures that were exposed to either 2% (blue) or 5% (purple) crude culture extract from symbiotic *Rhizopus microsporus* (RMsym). Incubation with solvent alone (DMSO) or apo-symbiotic *Rhizopus microsporus* (RMapo) has no effect on the viability of *P. aurantium*. Circles indicate independent replicated experiments (N = 3) ± one SEM (grey bars). One-way ANOVA with Tukey’s multiple comparison test (\**p<0.0001*, Supplementary file 3). **(B)** Photographs of yeast agar plates showing the predation plaque by *P. aurantium* (arrowheads). **(C)** HPLC profiles of crude extracts from symbiotic and endosymbiont-free *R. microsporus* showing a mixture of rhizoxin derivatives including the two major bacterial rhizoxin congeners (rhizoxin S1 and rhizoxin S2). The peak correlating to rhizoxin is marked with an *. Monitored at 310 nm. **(D)** Fluorescence microscopy images of symbiotic *R. microsporus* and endosymbiont-free *R. microsporus*. Green fluorescence indicates presence of endosymbionts (SYTO9). Scale bars: 5 μm.

### 2.3 The amoebicidal effect is delivered by rhizoxin

As *M. rhizoxinica* has a remarkable potential to produce toxic secondary metabolites, we investigated whether killing of the amoeba is caused by a bacterial metabolite. The fungal culture extracts were analysed for the presence of derivatives of the known cytotoxic macrolide rhizoxin (Partida-Martinez & Hertweck, 2005). HPLC profiles revealed a mixture of rhizoxin derivatives including the two major bacterial rhizoxin congeners (rhizoxin S1 and rhizoxin S2) in the symbiotic *R. microsporus* extract (Figure 3C) (Scherlach et al., 2006). The presence of *M. rhizoxinica* in the mycelium of symbiotic *R. microsporus* was confirmed by fluorescence microscopy (Figure 3D). In comparison, bacterial cells were absent in endosymbiont-free *R. microsporus* mycelium and none of the rhizoxin congeners were detected, suggesting that bacterial rhizoxins may cause lethal effects when ingested by *P. aurantium*.

To clarify whether the amoebicidal effect is delivered by a bacterial metabolite, we subjected *P. aurantium* to crude culture extracts from two representative endosymbiotic *Mycetohabitans* species, *M. rhizoxinica* (MR) and *Mycetohabitans endofungorum* (ME), which were isolated from their respective *R. microsporus* host (*R. microsporus* ATCC62417 and *R. microsporus* CBS112285, Figure 1) (Lackner, Mobius, et al., 2009; Partida-Martinez, Groth, et al., 2007). Both culture extracts from axenically grown bacteria (both 2% and 5%) kill all trophozoites of *P. aurantium* (Figure 4A,B). This effect is significant when compared to culture extracts from a rhizoxin-deficient *M. rhizoxinica* strain (*ΔrhiG*) and solvent controls (*p<0.0001*, Figure 4A,B—Supplementary file 4). HPLC analysis confirmed the presence of rhizoxin congeners in the two bacterial extracts, whereas none of the congeners are detected in extracts from the rhizoxin-deficient *M. rhizoxinica* strain (Figure supplement 3).

**Figure 4.**
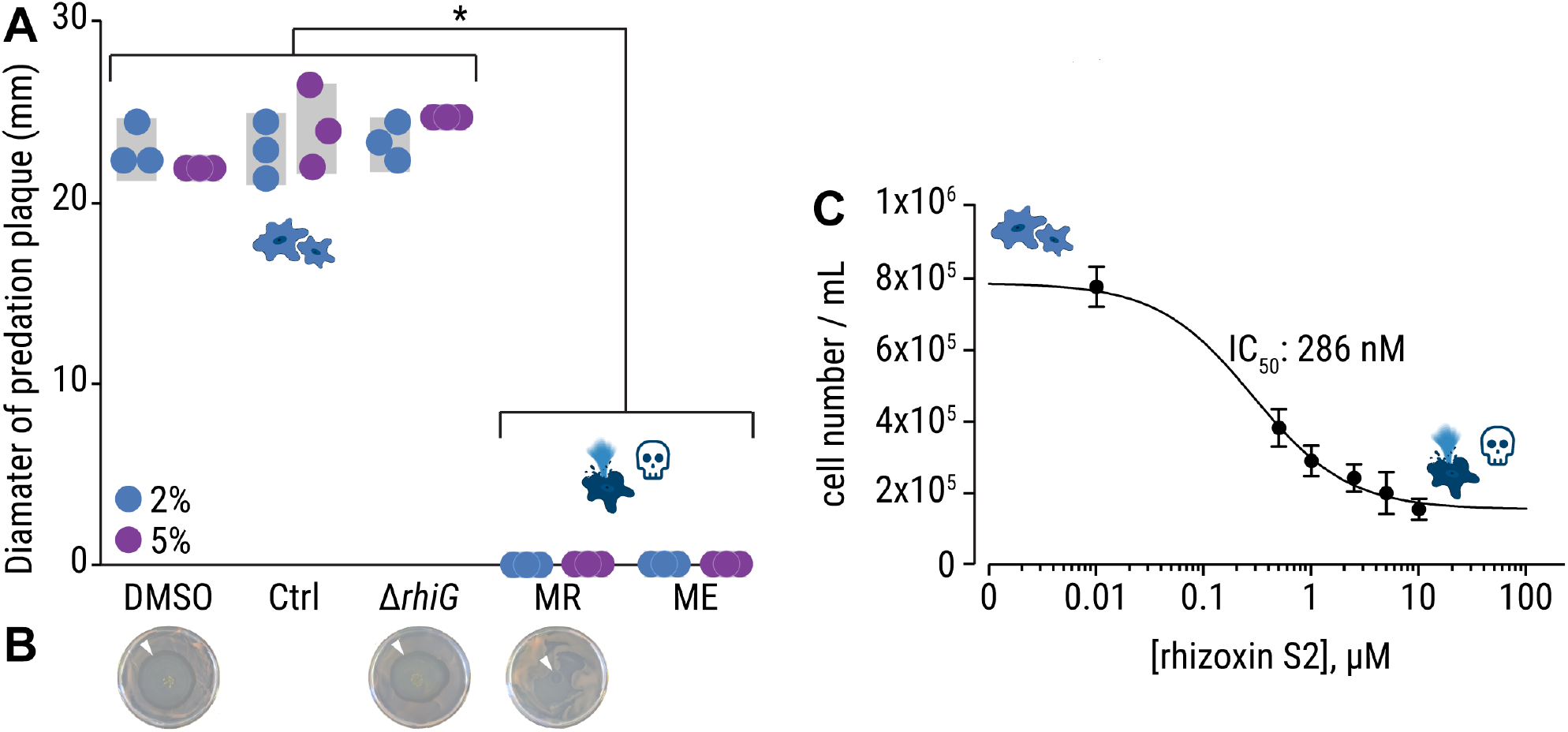
Culture extracts from axenic *Mycetohabitans* sp. kill *Protostelium aurantium*. **(A)** The viability of *P. aurantium*, indicated by the diameter of the predation plaque (clearance of yeast), is significantly reduced in cultures that were exposed to either 2% (blue) or 5% (purple) crude culture extract from axenically grown endosymbiotic *Mycetohabitans rhizoxinica* HKI-0454 (MR) or *Mycetohabitans endofungorum* HKI-0456 (ME). Incubation with solvent alone (DMSO), extract from culture medium (Ctrl), or rhizoxin-deficient *M. rhizoxinica (ΔrhiG*) has no effect on the viability of *P. aurantium*. Circles indicate independent replicated experiments (N = 3) ± one SEM (grey bars). One-way ANOVA with Tukey’s multiple comparison test (\**p<0.0001*, Supplementary file 4). **(B)** Photographs of yeast agar plates showing the predation plaque by *P. aurantium* (arrowheads). **(C)** Liquid toxicity assay of *P. aurantium* supplemented with the bacterial rhizoxin S2. Data points represent three independent replicated experiments (N = 3) ± one SEM.

To test whether bacterial rhizoxins kill *P. aurantium*, one of the major bacterial rhizoxin congeners (rhizoxin S2) was isolated from *M. rhizoxinica* as described previously (Scherlach et al., 2006) and its potency was assessed in a liquid *P. aurantium* toxicity assay. The sensitivity of *P. aurantium* (IC_50_ = 286 nM, 95% CI: 168–420 nM, Figure 4C) is comparable to the cytotoxic concentration against human HeLa cells previously reported for rhizoxin S2 (CC50 = 239 nM) (Scherlach et al., 2006). This confirms that bacterial rhizoxins are responsible for the killing of the fungivorous amoeba *P. aurantium*.

### 2.4 Endosymbionts protect *R. microsporus* from soil-dwelling nematodes

In addition to amoeba, fungi are also challenged by metazoan micropredators within the soil community. As nematodes are among the most abundant metazoan of the soil community (van den Hoogen et al., 2020), we investigated whether bacterial endosymbionts can protect *R. microsporus* from the ubiquitous soil nematode *C. elegans*, which has become a model system to study host-pathogen interactions (Smith et al., 2002).

*C. elegans*, co-cultured with *E. coli* OP50 as food source, was exposed to 2% culture extract in a liquid toxicity assay. The number of viable nematode worms in the suspension is directly related to the *E. coli* cell density (OD_600_). The positive control, 18 mM boric acid, kills the majority of worms (indicated by high OD_600_ values). As expected, the solvent control (DMSO) and extract from the *ΔrhiG* strain has no effect on nematode viability (indicated by low OD_600_ values). Fungal extracts of apo-symbiotic *R. microsporus* and bacterial extracts (*Mycetohabitans* sp. strains MR and ME) also show no effect on the survival rate of *C. elegans* with the majority of worms being alive (*p<0.0001*, Figure 5A—Supplementary file 5). However, a small difference in nematode viability was observed for the symbiotic *R. microsporus* extract. Significantly more worms die when compared to the extract from endosymbiont-free *R. microsporus (p<0.0002*, Figure 5A—Supplementary file 5), which is consistent with the effect of the fungal extracts on the survival of *P. aurantium* (Figure 2A).

**Figure 5.**
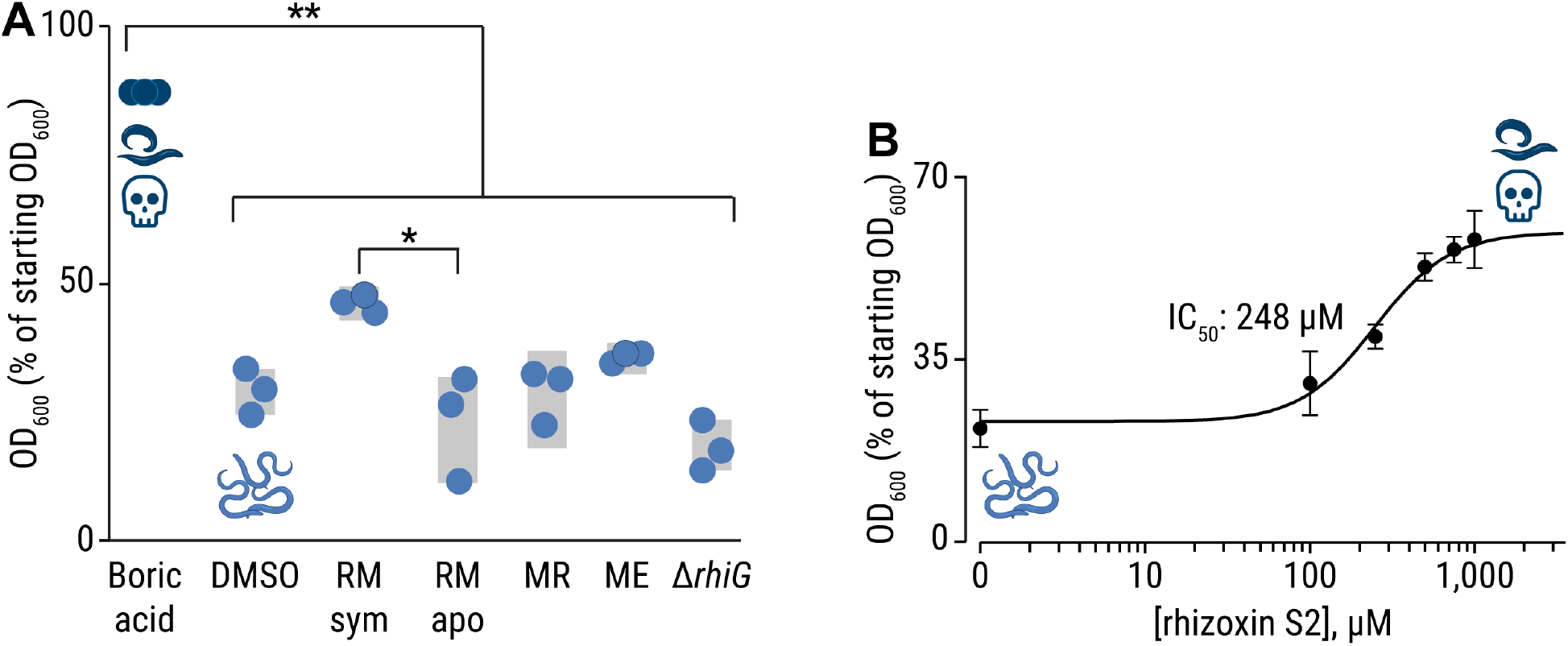
Toxicity effects of crude extracts and pure rhizoxin S2 on *Caenorhabditis elegans*. **(A)** *C. elegans*, co-incubated with *E. coli* OP50 cells as food source, were exposed to 2% crude culture extracts from symbiotic *Rhizopus microsporus* (RMsym), endosymbiont-free *Rhizopus microsporus* (RMapo), axenically grown endosymbiotic *Mycetohabitans rhizoxinica* HKI-0454 (MR), *Mycetohabitans endofungorum* HKI-0456 (ME), rhizoxin-deficient *M. rhizoxinica* (*ΔrhiG*). As the number of viable nematode worms in the suspension is directly related to the *E. coli* cell density, OD_600_ values were plotted as percent of the starting OD_600_. Incubation with 18 mM boric acid (positive control) kills most of the nematodes (*E. coli* density of 80%), while exposure to crude culture extracts does not affect *C. elegans* viability, which is comparable to the solvent control (DMSO). Circles indicate independent replicated experiments (n = 3) ± one SEM (grey bars). One-way ANOVA with Tukey’s multiple comparison test (ns: not significant, \*\**p<0.0001*, \**p<0.0002*, Supplementary file 5). **(B)** Liquid toxicity assay of *C. elegans* supplemented with the bacterial rhizoxin S2. Data points represent three independent replicated experiments (n = 3) ± one SEM.

As it was previously shown that the mobility of *C. elegans* decreases when directly in contact with *M. rhizoxinica* (Estrada-de Los Santos et al., 2018), we elucidated the IC_50_ for rhizoxin S2 in a dose-dependent toxicity assay (IC_50_ = 248 μM, 95% CI: 187–329 μM, Figure 5B). The inhibitory concentration against *C. elegans* exceeded the one found for *P. aurantium* (IC_50_ = 286 nM) by three orders of magnitude. This is in line with the observation that the majority of worms survived the treatment with crude culture extracts. The comparably low toxicity towards this nematode may be explained by the mode of rhizoxin delivery to *C. elegans* during exposure to extracts.

As the bacterivorous *C. elegans* does not feed on fungi in its natural environment, we explored the potential protective effect of fungal endosymbionts in an ecologically relevant setting. The interaction between the predatory nematode *Aphelenchus avenae* and *R. microsporus* was investigated. *A. avenae* was chosen as a model micropredator because of its ability to feed on fungal hyphae (Okada & Kadota, 2003). In addition, *A. avenae* and *R. microsporus* share the same ecological niche, as both species are globally distributed and inhabit temperate soils (Lackner, Mobius, et al., 2009; Okada & Kadota, 2003).

To study the feeding behaviour of *A. avenae*, nematodes were cultured on symbiotic *R. microsporus* as well as endosymbiont-free *R. microsporus* for 2–4 weeks. Following co-incubation, nematodes were harvested, and their viability was assessed by calculating their liveliness ratio (LR) as a measure of fitness. The LR was defined as the ratio of the area covered by a worm, divided by the area of the worm itself, and scaled to the full length of the time-lapse movie (30 sec). Nematodes with LR values below 1.4 were considered immobile, whereas a fast-moving worm would be characterized by a high LR value (the faster the worm’s movement, the higher the LR value). Nematodes grown on apo-symbiotic *R. microsporus* are healthy and active, as indicated by a LR of 6.24 ± 0.64, while worms living on symbiotic *R. microsporus* are significantly less lively (LR = 3.96 ± 0.57, *p<0.05*, Figure 6A,B—Supplementary file 6). Co-incubation of *A. avenae* with apo-symbiotic *R. microsporus* in a micro-channel slide confirmed active feeding of *A. avenae* on *R. microsporus* lacking endosymbionts (Figure 6C—video 1). *A. avenae* pierces the fungal cell wall with a stylet and feeds on the fungal cytoplasm by sucking (Schmieder et al., 2019). Sucking is facilitated by muscle contractions of the oesophagus, which is clearly visible in Video 1. After feeding, the nematode leaves a wound in the fungal cell wall, which leads to the release of fungal cytoplasm into the micro-channel (Figure 6C—video 1). In contrast, we did not observe active feeding on symbiotic *R. microsporus* hyphae, with the majority of the worms being dead (Figure supplement 5—video supplement 2). As indicated by our *C. elegans* data, the protective effect against *A. avenae* is likely mediated through the secondary metabolite rhizoxin, which is produced by endosymbionts living with in the fungal hyphae (Figure 7). In line with this model, *A. avenae* is likely rhizoxin sensitive, as the β-tubulin amino acid sequence of the closely related fungivorous nematode *Bursaphelenchus okinawaensis* harbours asparagine at amino acid position 100 (Figure supplement 4). These results highlight a defensive function of an endofungal symbiotic bacterium against protozoan and metazoan predators.

**Figure 6.**
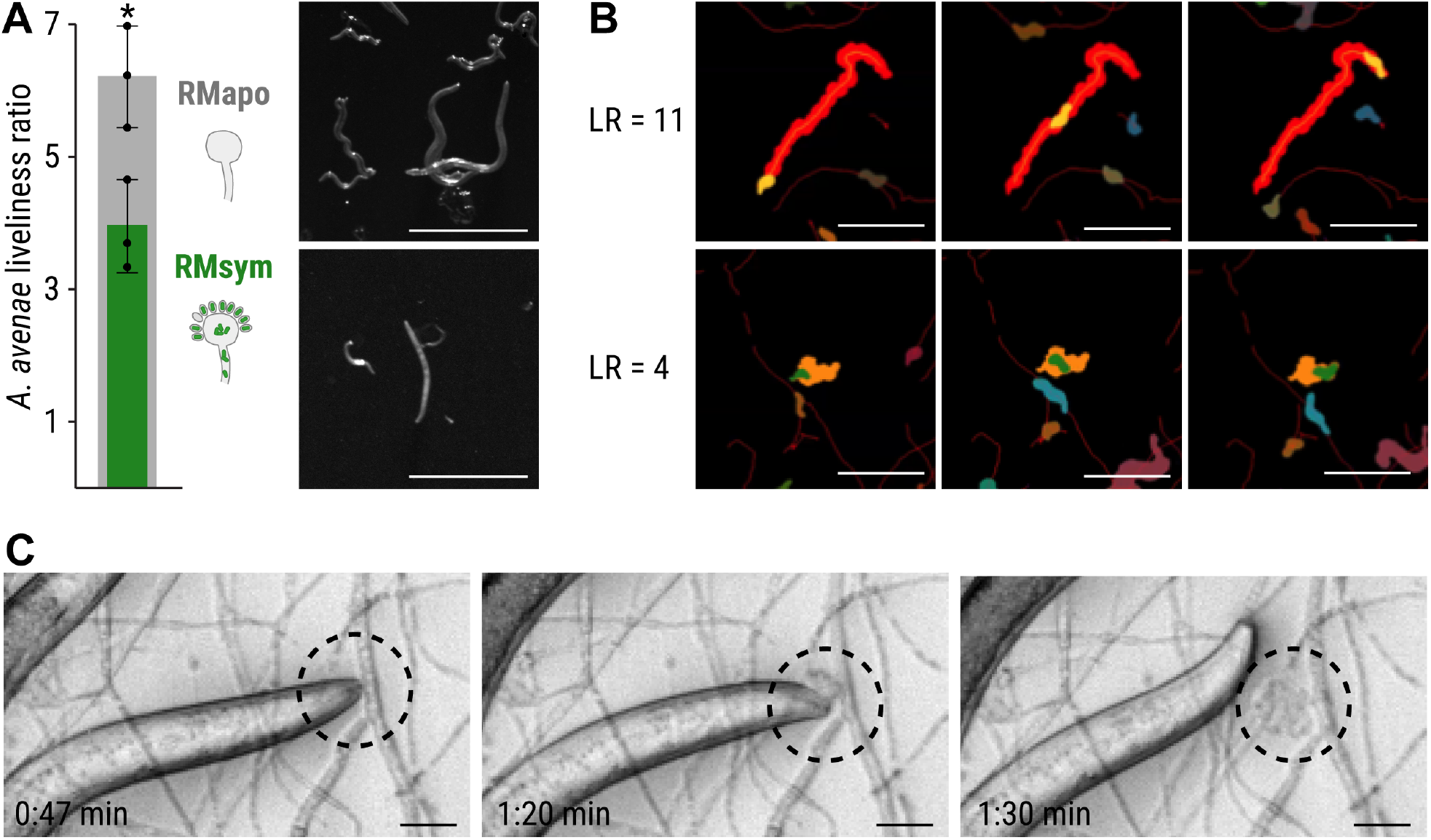
Feeding inhibition of *Aphelenchus avenae* on *Rhizopus microsporus*. **(A)** *A. avenae* was coincubated with symbiotic *R. microsporus* (RMsym) or endosymbiont-free *R. microsporus* (RMapo) for 2–3 weeks. Nematode movement was recorded using a stereomicroscope with a frame rate of 1 fps. The liveliness of the worms was calculated from the ratio of the area covered by a worm, divided by the area of the worm itself, and scaled to the full length of the movie. The minimum scaled liveliness ratio (LR) for a live worm was set to 1.5, below this value the worm was declared inactive/dead. N = 3 independent replicated experiments ± one SEM. Unpaired *t*-test with Welch’s correction (\**p<0.05*, Supplementary file 6). Microscope images of *A. avenae* used for analysis. Scale bars: 500 μm. **(B)** Illustrations of the LR at high (upper) and medium (lower) values. Left: the worm shown in orange covers the red footprint area during the time course of the experiment. Here are shown images from the first (top row), middle (middle row) and final (bottom row) time points of the movie. The activity of a worm was characterized by dividing the endpoint footprint by the area of the worm at each time point. The resulting LR was 11.5 for the worm in the left column, thus indicating a very active nematode. Right: A less active worm (green area) covered a smaller footprint (orange area) as shown by the LR value of 4.0. Scale bars: 300 μm. See the live videos of the segmented worms and their footprints in Video supplement 6 and Video supplement 7 for the worms of LR = 11.5 and LR = 4.0, respectively. **(C)** Time-lapse images of *A. avenae* feeding on endosymbiont-free *R. microsporus* (black circle). Endosymbiont-free *R. microsporus* ATCC62417/S was co-incubated with *A. avenae* for 24 hrs in a micro-channel slide (Ibidi) and feeding was recorded on a spinning disc microscope (Video 1). Scale bars: 20 μm. No feeding was observed in worms that were co-incubated with symbiotic *R. microsporus* (Figure supplement 5—video supplement 2).

**Figure 7.**
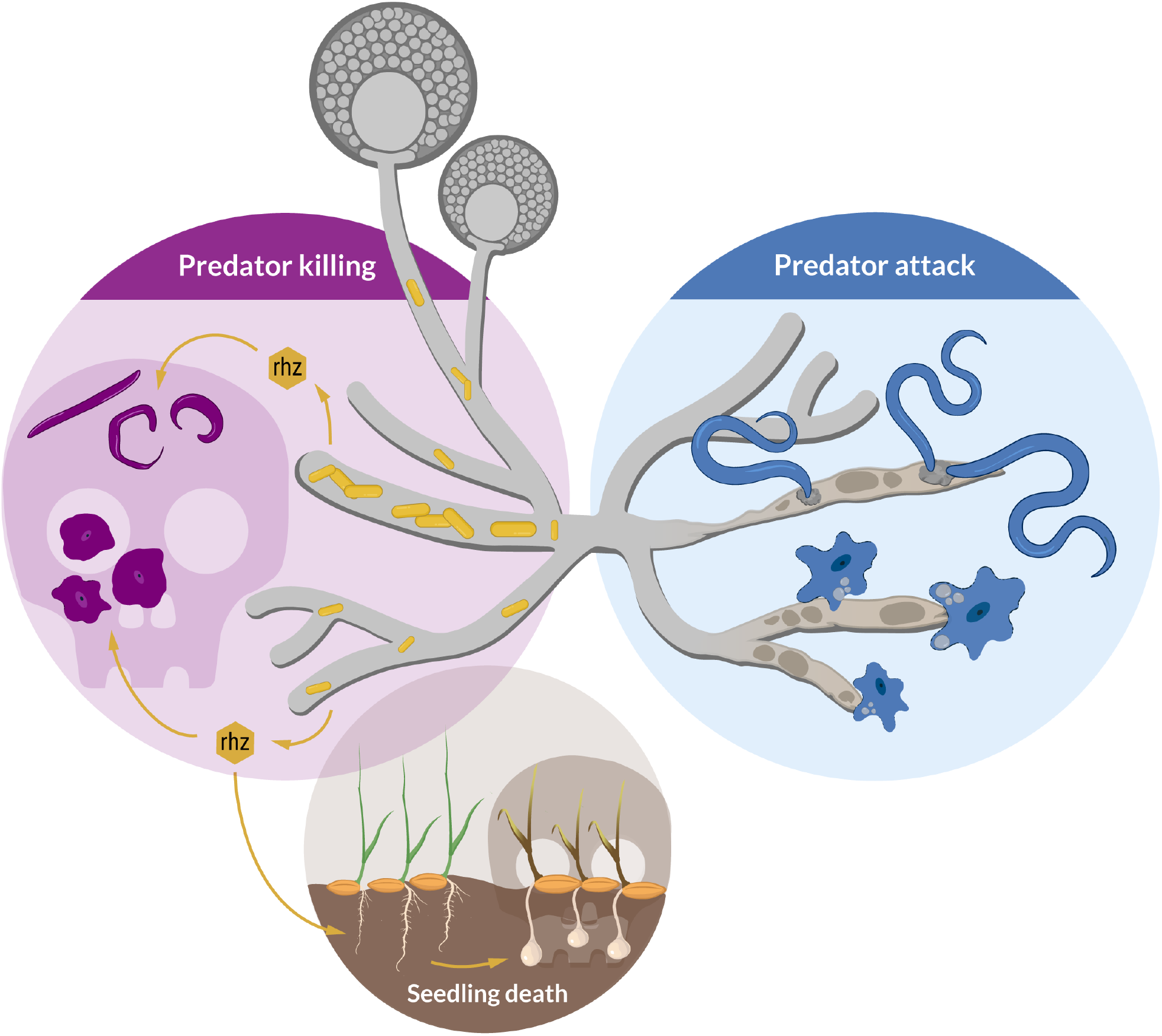
Schematic model of the ecological role of rhizoxin-producing endofungal bacteria (*Mycetohabitans rhizoxinica*). The fungal host (*Rhizopus microsporus*) utilises the bacterial secondary metabolite rhizoxin to fend off fungivorous micropredators such as amoeba and nematodes. Absence of endofungal bacteria leads to *R. microsporus* being attacked and subsequently killed by protozoan and metazoan predators. The establishment of the *Rhizopus* – *Mycetohabitans* symbiosis may have first developed to provide protection against fungal predators with the emergence of plant pathogenicity developing later.

## 3 Discussion

Prey-predator interaction is a major driver of biodiversity (Brodersen et al., 2018). Within the soil microbiome, fungi are constantly threatened by fungivorous organisms selecting for strategies to defend themselves against predators (Künzler, 2018; Li et al., 2007; Spiteller, 2015). In this study we revealed that endosymbionts protect the phytopathogenic fungus *R. microsporus* against fungivorous protozoan and metazoan predators. Using a combination of co-culture experiments, cytotoxicity assays, and fluorescence microscopy we report an amoebicidal and nematicidal effect of rhizoxin, a secondary metabolite produced by the bacterium *M. rhizoxinica* residing within the hyphae of *R. microsporus* (Scherlach et al., 2012).

Fungivorous amoebae are ubiquitous in soil and leaf litter with *P. aurantium* being a prime example of a species that feeds on a wide range of unicellular yeasts as well as conidia and hyphae of filamentous fungi (Hillmann et al., 2018; Radosa et al., 2019). We observed that *P. aurantium* also ingests spores of *R. microsporus* via phagocytosis as shown by fluorescence microscopy. However, ingestion of spores has lethal consequences for the amoeba when *R. microsporus* spores contain the toxin-producing endosymbiont (Partida-Martinez & Hertweck, 2005). While it was previously suggested that *Ralstonia pickettii*, an endosymbiont of *R. microsporus*, secretes growth suppressing factors against the soil-dwelling amoeba *Dictyostelium discoideum* (Itabangi et al., 2020), we present the first report of rhizoxin-mediated killing of a fungivorous amoeba. These results highlight that endosymbionts protect their fungal host from being attacked by mycophagous amoeba and it is the first report identifying a Mucoromycota fungus as food source of *P. aurantium*. Rhizoxin toxicity is most likely not limited to *P. aurantium*, but may have a rather wide biological range among the kingdom of Amoebozoa, as similar effects have been observed for the distantly related parasitic amoeba *Entamoeba histolytica*, whose survival rate was reduced by 58% when exposed to methanol extracts (1 g/L) from *R. microsporus* cultures (Kapilan & Anpalagan, 2015).

Rhizoxin efficiently binds to β-tubulin, leading to the potent depolarisation of microtubules and subsequent mitotic arrest in humans and plants (Prota et al., 2014). A conserved residue at amino acid position 100 of the β-tubulin protein is important for rhizoxin binding and subsequently rhizoxin sensitivity (Schmitt et al., 2008). In line with these observations, the *P. aurantium* β-tubulin protein harbours the amino acid asparagine at position 100 (Figure supplement 4), which is an important feature in rhizoxin-sensitive fungi (Schmitt et al., 2008). Depolarisation of microtubule in amoeba has severe consequences, as β-tubulins have been shown to be essential parts of the microtubule network in *D. discoideum*, important for cell polarity, migration and the movement of intracellular particles (Triviños-Lagos et al., 1993). Intriguingly*, P. aurantium* requires substantial condensation of actin filaments when grabbing and invading the rather large fungal hyphae (Radosa et al., 2019). It is thus conceivable that this part of the cytoskeleton, which actually enters fungal cells would be a primary and highly effective target of the intrafungal rhizoxin.

We further uncovered that rhizoxin can also protect against higher eukaryotic predators from the kingdom of Metazoa. Cytotoxicity assays revealed a lethal effect of this compound on the soil-dwelling model nematode *C. elegans*. The inhibitory concentration is far higher when compared to *P. aurantium*. This relatively low sensitivity towards rhizoxin S2 might explain why crude culture extracts showed a minor effect on *C. elegans* viability. However, it is well known that the potencies of different rhizoxin congeners can vary greatly between organisms. For example, picomolar concentrations of rhizoxin S2 inhibit proliferation of leukaemia cell lines (GI50 1.6 pM), while growth inhibition of the plant pathogenic fungus *Phytophthora ramorum* requires rhizoxin concentrations in the micromolar range (Loper et al., 2008; Scherlach et al., 2006). Thus, the effectiveness against *C. elegans* fits within the range of potencies reported for rhizoxin and may greatly depended on the biological setting. While *P. aurantium* and *A. avenae* actively feed on the fungal cytoplasm, *C. elegans* is bacterivorous and thus, unlikely to be directly exposed to toxins from fungal endosymbionts. However, the alignment of all six *C. elegans* β-tubulin amino acid sequences suggests that also *C. elegans* is sensitive to rhizoxin due to the presence of asparagine at amino acid position 100 (Figure supplement 4) (Schmitt et al., 2008). Since worm tubulins play an essential role during all phases of the cell and life cycle (Hurd, 2018), it is likely that rhizoxin-induced killing of *C. elegans* is mediated through microtubule depolarisation.

The combination of stereomicroscopy, automated image analyses, and quantification of nematode movement demonstrated that the fungivorous nematode *A. avenae* actively feeds on *R. microsporus* that is lacking endosymbionts. In contrast, presence of the bacterial endosymbionts in symbiotic *R. microsporus* causes death in the majority of worms. This lethal effect is either due to nematode starvation or active feeding on *R. microsporus* and subsequent ingestion of toxic compounds. However, since *C. elegans* does not feed on axenic *M. rhizoxinica* leading to nematode death by starvation (Estrada-de Los Santos et al., 2018) it is likely that the presence of *Mycetohabitans* sp. inside the fungal hyphae protects *R. microsporus* from predatory nematodes. These results highlight a defensive function of an endofungal symbiotic bacterium against a metazoan predator and may explain the very limited *A. avenae* range of prey among Mucoromycota fungi with only two species (*Mucor hiemalis* and *Mortierella verticillata*) known to act as a food source (Büttner et al., 2021; Ruess et al., 2000).

By discovering fungivorous predation on *R. microsporus* this study adds another dimension to the tripartite interaction between fungal host, symbiont, and rice plants (Scherlach et al., 2013) and opens an interesting evolutionary perspective. The establishment of the *Rhizopus* – *Mycetohabitans* symbiosis may have originally developed to provide protection against fungal predators and only later facilitated the emergence of plant pathogenicity. The interactions between Mucoromycota and their bacterial endosymbionts are an ancient phenomenon dating back as far as 400 million years (Mondo et al., 2012). Flowering land plants like rice developed far later (134 million years ago) than protozoan and metazoan predators such as amoebae and nematodes (400 million years ago) (Poinar, 2011; Strullu-Derrien et al., 2019). Thus, the establishment of a mutualistic interaction between *Rhizopus* and rhizoxin-producing *Mycetohabitans* (Lastovetsky et al., 2016) may have allowed the fungal host to evade predator attack thereby gaining an evolutionary advantage over aposymbiotic or rhizoxin-negative symbiotic fungi.

Here, we uncovered an unexpected role of rhizoxin. In addition to causing blight disease in rice seedlings, we show that this bacterial secondary metabolite is utilised by the fungal host to successfully fend off micropredators (Figure 7). This anti-predator effect of toxin-producing endofungal bacteria of *Rhizopus* is an important addition to a similar observation in *Mortierella* species (Büttner et al., 2021) and points to a more widespread ecological role of endosymbionts. In line with this model, endosymbiont-containing *Rhizopus* species are globally distributed and the toxin produced by the endosymbionts is lethal to most eukaryotes, including insects, vertebrates, and fungi. It is thus conceivable that animal predation represents an evolutionary driving force to maintain endosymbionts in non-pathogenic fungi.

## 4 Material and methods

### Strains and growth conditions

Liquid cultures of *Protostelium aurantium var. fungivorum* were grown to confluency in 2 mM phosphate buffer (pH 6.2; PB) supplemented with *Rhodotorula mucilaginosa* as a food source in standard-sized Petri dishes at 22 °C (Hillmann et al., 2018).

The endobacteria *Mycetohabitans rhizoxinica* HKI-0454 (MR) and *Mycetohabitans endofungorum* HKI-0456 (ME) were isolated from the mycelium of *Rhizopus microsporus* ATCC62417 and *Rhizopus microsporus* CBS112285, respectively (Lackner, Mobius, et al., 2009; Partida-Martinez, Groth, et al., 2007). Axenic *Mycetohabitans* sp. cultures were maintained at 30 °C in MGY M9 medium (10 g/L glycerol, 1.25 g/L yeast extract, M9 salts) or Standard I Nutrient Agar (Merck, Darmstadt, Germany) supplemented with 1% glycerol. Symbiotic *R. microsporus* ATCC62417 (RMsym) was treated with antibiotics to eliminate its endosymbionts (Partida-Martinez & Hertweck, 2007) resulting in the apo-symbiotic fungal strain ATCC62417/S (RMapo). As the lack of endosymbionts also abolishes the production of rhizoxin, we confirmed the absence of symbionts from ATCC62417/S by checking for rhizoxin congeners in ATCC62417/S culture extracts via HPLC (Scherlach et al., 2006). Both *R. microsporus* strains (ATCC62417 and ATCC62417/S) were cultivated on Potato Dextrose Agar (PDA; Becton, Dickinson & Company, Sparks, MD, USA) at 30 °C.

*Caenorhabditis elegans* wild-type N2 (var. Bristol), purchased from the *C. elegans* Genetics Centre (CGC, University of Minnesota, USA), was grown and maintained on nematode growth medium (NGM) containing *E. coli* OP50 as food source (Stiernagle, 2006).

### *Protostelium aurantium* amoeba predation assay on *Rhizopus microsporus* spores

*R. microsporus* was grown on PDA plates for 7 days. Spores were harvested using 12 mL NaCl (0.15 M). The spore solution was centrifuged at 9,000 rpm for 15 min. The resulting pellet was resuspended in 1 mL 50% glycerol and spores were counted in a Thoma Chamber.

A total of 10^5^ spores of *R. microspores* were seeded in 96-well tissue culture plates containing 200 μL Czapek-Dox medium (CZD, Sigma-Aldrich Chemie, Munich, Germany). These spores were confronted with *P. aurantium* directly (dormant spores) or after pre-incubation at 30 °C for 3 hrs (swollen spores), in a prey-predator ratio of 10:1. Trophozoites of *P. aurantium* were pre-grown in PB with *R. mucilaginosa* as a food source. To measure the metabolic activity of the surviving fungus, 0.002% [*w*/*v*] resazurin (Sigma-Aldrich, Taufkirchen, Germany) was added to the same wells. After the incubation period (20 hrs at 22 °C), the fluorescent intensity of resorufin was measured using an Infinite M200 Pro fluorescence plate reader (Tecan, Männedorf, Switzerland). Survival rates were determined by the difference in the metabolic activity of the fungus after amoeba confrontation and amoeba-free controls.

To visualize predation of *P. aurantium* on spores of *R. microsporus*, approximately 500 μL of spore suspension was mixed with FITC staining solution (1 mg/10 mL 0.1 M Na_2_CO_3_) and incubated at 37 °C for 30 min and 8000 rpm, under light exclusion. Following incubation, the spores were centrifuged and washed three times with PB to remove unbound staining. Stained spores were co-incubated with *P. aurantium* trophozoites and visualized using a fluorescence spinning disc microscope (Axio Observer microscope-platform equipped with Cell Observer SD, Zeiss) with an excitation/emission range of 495/519 nm.

### Culture extraction and compound isolation

Axenic *Mycetohabitans* sp. cultures were grown in 400 mL MGY containing 10% TSB (17 g/L tryptone, 3 g/L soy, 5 g/L NaCl, 2.5 g/L K_2_HPO_4_, 2.5 g/L glucose) in 1-L baffled Erlenmeyer flasks for 7 days at 30 °C and 110 rpm. Fungal strains (ATCC62417 and ATCC62417/S) were grown on 20 PDA-containing standard-sized Petri dishes at 30 °C for 7 days. Both liquid bacterial cultures and fungal agar plates were exhaustively extracted with 400 mL ethyl acetate. Extracts were concentrated on a rotary vacuum evaporator and then dried. Dry extracts were dissolved in 1 mL methanol and analysed via HPLC as described previously (Scherlach et al., 2012). Following HPLC analysis, crude extracts were dried and dissolved in 1 mL DMSO for bioactivity assays. The volume of DMSO was adjusted for the bacterial extracts making all samples equal to a starting OD_600_ of approximately 3.5.

For the isolation of rhizoxin S2, *M. rhizoxinica* HKI-0454 was cultured in 5.6 L MGY 10% TSB equally disturbed across 1-L baffled Erlenmeyer flasks. Compounds were isolated as described previously (Scherlach et al., 2012; Scherlach et al., 2006).

### *Protostelium aurantium* plaque assay

Amoeba cells were seeded in the 96-well plates (Costar^®^, TC-treated; Corning, NY, US) at a concentration of 10^5^ cells/mL in PB. Cells were either left untreated or incubated in the presence of 2% or 5% of crude extract from bacterial or fungal cultures for 1 hr. Incubation with media or solvent (DMSO) was included as controls. Afterwards, 20 μL of cell suspension was pipetted in the middle of a PB agar plate covered with a dense layer of *R. mucilaginosa* as a food source. The predation plaque, appearing as a halo due to the clearance of the yeast, was measured after five days. Each experiment was performed in three biological replicates.

### *Protostelium aurantium* cytotoxicity assay

Amoeba cells were seed in 24-well plates (Falcon^®^, TC-treated, Corning, NY, US), at a concentration of 10^6^ cells/mL in PB. Rhizoxin S2 was prepared in the working concentration of 100 μM. In a defined concentration range, rhizoxin S2 was added to the cells and filled up to final volume of 500 μL. Amoeba cells were further incubated in the presence of *R. mucilaginosa* for 24 hrs. The next day, cells were scraped off the bottom of the wells and the number of viable cells was counted manually in an improved Neubauer counting chamber. Each experiment was performed in three biological replicates. A four-parameter sigmoidal concentration-dependent response curve was fitted using GraphPad Prism Version 6.03 (GraphPad Software, La Jolla, California, USA, www.graphpad.com) and the 50% inhibitory concentration (IC_50_) and 95% confidence intervals (CI) were determined.

### *Caenorhabditis elegans* liquid toxicity assay

For toxicity assays, *C. elegans* was cultured on NGM OP50 agar plates at 20 °C for 4 days. Nematodes were harvested by washing the agar plates with 12 mL sterile K-medium (3.1 g/L NaCl, 2.4 g/L KCl). Worms were allowed to settle to the bottom through incubation at 4 °C for 20 min. The supernatant was carefully removed, and the worms were washed with 12 mL K-medium twice. On the last washing step, the supernatant was removed, and worms were resuspended in 5 mL fresh K-medium.

Liquid OP50 cultures were grown in Lysogeny broth (10 g/L tryptone, 5 g/L yeast extract, 10 g/L NaCl, pH 7.0) at 37 °C overnight. Cells were harvested by centrifugation at 8,000 rpm for 5 min, resuspended in K-medium and the OD_600_ adjusted to 1.2. Individual wells of six-well cell culture plates (Costar^®^, Corning, NY, USA) were seeded with 1.76 mL *E. coli* suspension (OD_600_ of 1.2) and 200 μL of nematode suspension.

Crude culture extracts, dissolved in DMSO, were added to the wells (40 μL). DMSO served as blank control and 40 μL of 900 mM boric acid (final concentration of 18 mM) was used as positive control. To determine the natural viability of *E. coli* OP50 cells, wells without nematodes were included in each assay. All plates were incubated for 7 days at 20 °C, 90 rpm and the OD_600_ was measured every 24 hrs. The number of viable nematode worms in the suspension is directly related to the *E. coli* cell density. Mean OD_600_ values from three independent experiments (N = 3 biological replicates) were plotted as percent of the starting OD_600_ ± one SEM.

In the cytotoxicity assay, rhizoxin S2 was dissolved in DMSO and added (40 μL) to the wells with a final concentration range of 0.1–1000 μM (3 replicates per concentration). OD_600_ values were measured as described above and the IC_50_ value and 95% CI were calculated using GraphPad.

### *Aphelenchus avenae – Rhizopus microsporus* co-culture

The nematode *Aphelenchus avenae* (Bastian, 1865) was kindly provided by Dr. Markus Künzler (ETH Zürich, Switzerland). *A. avenae* was maintained on a non-sporulating strain of *Botrytis cinerea* (BC-3) growing on malt extract agar plates (MEA) supplemented with 100 μg/mL chloramphenicol at 20 °C (Kumar et al., 2007). For feeding assays, nematodes were harvested from co-cultures by Baermann funnelling overnight (Hooper et al., 2005; Walker & Wilson, 1960). Nematodes were collected in 50 mL falcon tubes and incubated at 4 °C for 2 hrs. The supernatant was removed, and worms were resuspended in 50 mL sterile K-medium supplemented with 25 μM kanamycin and 100 μM geneticin. The worm suspension was incubated at room temperature for 2 hrs to eliminate remaining fungal spores. Worms were washed with 50 mL K-medium twice, resuspended in 5 mL K-medium, and added (500 μL) to PDA plates containing 7-day old cultures of symbiotic *R. microsporus* (ATCC62417) and apo-symbiotic *R. microsporus* (ATCC62417/S) in triplicates. After incubation at 21 °C for 2 – 4 weeks, worms were harvested from co-cultures by Baermann funnelling as described above. Following antibiotic treatment and washing with K-medium, worms were transferred on to sterilisation agar plates (15 g/L agar, 50 μg/mL kanamycin, 200 μM geneticin) and incubated at 21 °C overnight (Bleuler-Martínez et al., 2011). Nematode movement was recorded in timelapse videos (1 min) with a frame rate of 1 fps using a Zeiss Axio Zoom.V16 Stereomicroscope (Zeiss, Oberkochen, Germany).

### Image analysis and mathematical quantification of *Aphelenchus avenae* migration

Transmitted-light time series images of free-moving worms were analysed via an automated image processing and quantification algorithm. The raw images were provided in the native Carl-Zeiss image format “CZI”, whereas the processing was carried out in a novel graphical image analysis language JIPipe (www.jipipe.org), available as a plugin in ImageJ (v.1.53c). Images recorded both at 25× and 32× magnification were used, without having to modify the analysis steps or controlling parameters. At these magnifications individual worms were easily identifiable without fluorescence labelling, thus avoiding interference with biological function (Cseresnyes et al., 2020). The worm number in the microscope’s field of view was high enough to provide statistically meaningful quantification results. The step-by-step workflow is summarized in Figure supplement 6, whereas images depicting representative intermediate results of the processing are presented in Figure supplement 7. The images were first corrected for uneven transmitted light illumination, followed by Laplacian sharpening (3 × 3 pixels) and a Hessian filter with 5-pixel kernel diameter (Cseresnyes et al., 2018). Here the largest Hessian eigenvalues were used to build an image that showed the worms at enhanced contrast due to the continuously curved body outline of a typical nematode. The Hessian eigenimage was further processed to produce a high-fidelity segmentation of individual worms: a Gaussian blurring using a 3-pixel disk structural element was followed by thresholding using the Default algorithm (i.e. a modified IsoData algorithm) and morphological dilation (1 pixel) and hole filling. The non-worm image elements were removed by applying the Remove Outliers command at sizes below 50 pixels. The per-image segmentation of individual worms was followed by tracking using the connected components algorithm of JIPipe (“Split into connected components” node). The tracking workflow was carried out by two parallel branches: one algorithm extracted the per-worm and per-track information of individual worm areas, whereas the other branch calculated the total area covered by each worm whilst counting each worm-covered pixel only once. The latter workflow thus provided a footprint of a worm covered during the time-series experiment (Figure supplement 8—video supplement 5). In the next JIPipe compartment, the footprint of the worm was divided by the individual worm areas recorded at each time point; the resulting ratio was 1.0 for a fully immobile worm, whereas more active worms produced higher ratios. This ratio was termed Liveliness Ratio (LR), because it quantifies the agility of a worm. The average ratio per track was calculated by taking the arithmetic mean and standard deviation of the per time—point ratio values. The ratio varied with time due to variance of the apparent area of the worm as measured by the per-image segmentation algorithm. Those tracks that were characterized by very high standard deviations were excluded from further analysis, because a high standard deviation would indicate that the segmentation was unreliable at some timepoint(s) of the track, e.g. due to a merging of two or more worms into a cluster that could not be resolved by the segmentation algorithm. This ratio was called Liveliness Ratio, or LR, because of its description of the agility of a worm. Nematodes with LR values below 1.4 were considered immobile, whereas a fast-moving worm would be characterized by a high LR value with an average of 5.5 ± 0.5 over all experiments and conditions. The average and standard deviation LR results were saved in CSV file formats per experiment and condition for further analysis. An example of the time series of segmented worms and their tracks can be viewed in Video supplement 3. Video supplement 4 shows the velocity distribution of the same movie. The isotropic nature of the velocity vector-distribution points to a random walk-type worm movement in the movie.

### *Aphelenchus avenae* feeding on *Rhizopus microsporus*

To monitor feeding of *A. avenae* on *R. microsporus*, a 0.2 μm Luer μ-slide (Ibidi GmbH, Gräfelfing, Germany) was filled with sterile Potato Dextrose Broth (Becton, Dickinson & Company). *R. microsporus* mycelium (ATCC62417 or ATCC62417/S) was added to one of the inoculation holes and slides were incubated at 30 °C for two days to allow hyphae to grow into the micro-channel. Sterilised *A. avenae* (see above) were added to the second inoculation hole and slides were incubated at 21 °C overnight. Feeding of *A. avenae* on *R. microsporus* hyphae was observed using a Zeiss spinning disc microscope (Axio Observer microscope-platform equipped with Cell Observer SD, Zeiss).

### Statistical analysis

Raw data from *P. aurantium, C. elegans*, or *A. avenae* survival experiments were processed in MS Excel and statistical analysis was performed in GraphPad Prism 6.03. An unpaired *t*-test with Welch’s correction was used to study the following relationships: (i) ingestion of swollen *R. microsporus* spores vs. dormant spores by *P. aurantium;* and (ii) liveliness ratio of *A. avenae* feeding on either symbiotic *R. microsporus* or endosymbiont-free *R. microsporus*. To study the effect of fungal (symbiotic and apo-symbiotic *R. microsporus*) and bacterial (*Mycetohabitans* sp.) culture extracts on the survival of *P. aurantium* and *C. elegans*, one-way analysis of variance (ANOVA) was used in combination with the Tukey HSD test function. The Brown-Forsythe test was used to test for equal variance. For all statistical tests performed, p-values with *p<0.05* were considered statistically significant.

### Alignment of β-tubulin proteins

Amino acid sequences from β-tubulin genes of *Protostelium aurantium, Dictyostelium discoideum, Acanthamoebae castellanii, Caenorhabditis elegans*, and *Rhizopus oryzae* were downloaded from the Universal Protein Resource (https://www.uniprot.org/). β-tubulin amino acid sequences from *Rhizopus microsporus var. microsporus* and *Bursaphelenchus okinawaensis* were downloaded from GenBank. Sequence alignments were carried using ClustalW (Thompson et al., 1994). Alignments were generated using a gap open penalty of 10 and a gap extension penalty of 0.1 as implemented in the MEGA7 package (Molecular Evolutionary Genetics Analysis software, version 5.0) (Kumar et al., 2016).

## 5 Acknowledgments

*Caenorhabditis elegans* was provided by the CGC, which is funded by NIH Office of Research Infrastructure Programs (P40 OD010440). We thank Dr. Kirstin Scherlach for helpful advice and Prof. Dr. M. Künzler (ETH Zürich) for kindly providing *Aphelenchus avenae*. I.R. is grateful for financial support from the European Union’s Horizon 2020 Research and Innovation Programme under the Marie Skłodowska-Curie grant agreement No. 794343. Financial support by the Deutsche Forschungsgemeinschaft (DFG; German Research Foundation) by SFB 1127/2 ChemBioSys - 239748522 (to H.B. and C.H.) and Leibniz Award (to C.H.) is gratefully acknowledged. S.R. and F.H. were supported by the DFG under Germany’s Excellence Strategy – EXC 2051 – Project-ID 390713860. Z.C. and M.T.F. were funded by the DFG – project number 316213987 – SFB 1278 “PolyTarget” (Z01). R.G. was funded by the International Leibniz Research School for Microbial and Biomolecular Interactions Jena – ILRS Jena. Figures were created with biorender.com.

## 6 Additional Information

### Funding

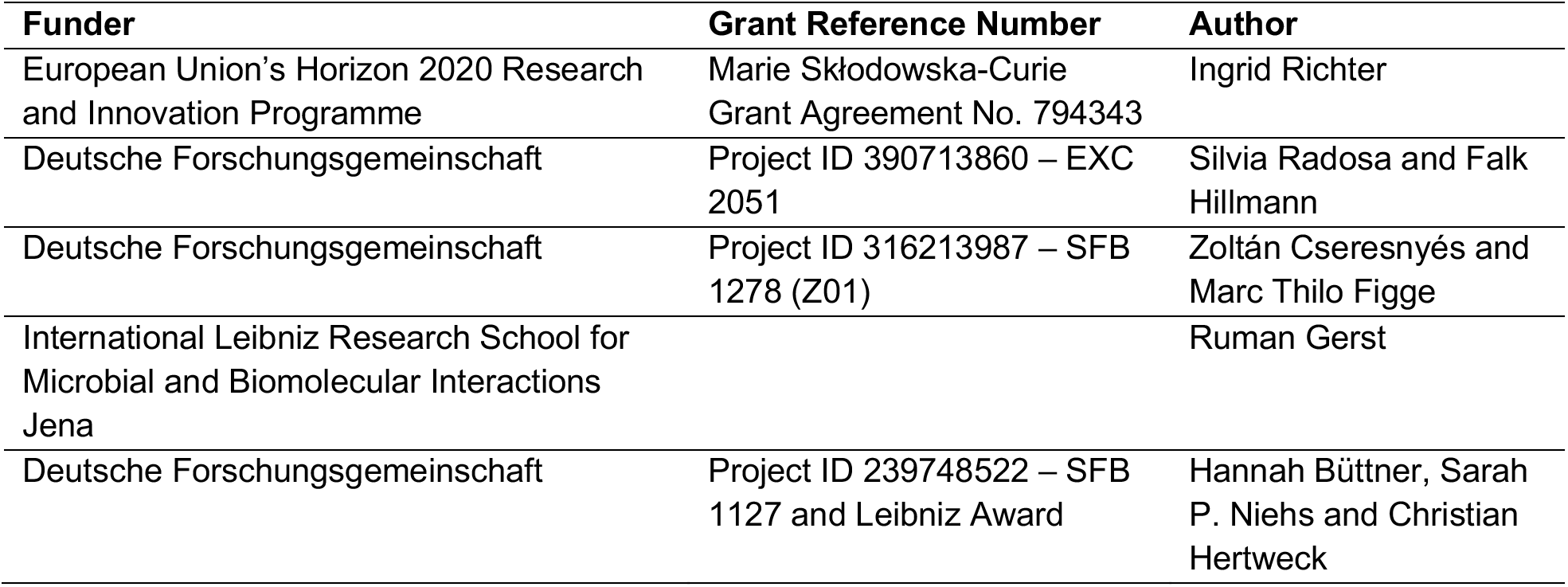

## Author contributions

IR: project design, planning and performing nematode assays, performing culture extractions; analysis and interpretation of data, writing original draft; SR: project design, planning and performing amoeba assays; drafting or revising the article; ZC: conducting nematode movement analysis; drafting or revising the article; IF: planning and performing amoeba assays; HB: supported nematode assays; SPN: supported compound isolation; RG: conducting nematode movement analysis; MTF: guiding computational image analysis; FH: project design, drafting and revising the article; CH: project design, drafting and revising the article.

## Data availability

All data generated or analysed during this study are included in the manuscript and in the supporting files. Source data files have been provided for Supplemental videos.

## File captions

**Video 1. *Aphelenchus avenae* feeding on endosymbiont-free *Rhizopus microspores*.** Endosymbiont-free *R. microsporus* ATCC62417/S was co-incubated with *A. avenae* for 24 hrs in a micro-channel slide (Ibidi) and feeding was recorded on a spinning disc microscope.

**Video supplement 1. Predation of *Protostelium aurantium* on swollen spores from *Rhizopus microsporus*.** Time-lapse movie showing ingestion of a swollen *R. microsporus* spore (stained with FITC) by *P. aurantium*. Scale bar: 5 μm.

**Video supplement 2. *Aphelenchus avenae* co-incubated with symbiotic *Rhizopus microsporus***. R. microsporus ATCC62417 was co-incubated with *A. avenae* for 24 hrs in a micro-channel slide (Ibidi). Timelapse movie, recorded on a spinning disc microscope, showing dead/unhealthy nematodes.

**Video supplement 3. The segmented worms and their tracks of a time series experiment.** The worms and the tracks are shown here as provided by the automated tracking algorithm applied to a time series experiment. The worms are coloured randomly, whereas the tracks (thin lines) are coloured from blue to red for each track, blue corresponding to time zero and red to the final time point. When worms merge, they become of the same colour until they separate again.

**Video supplement 4. The X component of the per-worm and per time-point velocity vector of each worm as a function of the Y component of the velocity vector.** The time series shows the velocity vector components at individual time points, playing from time zero to the final time point. Line colours correspond to the time, whereas the worm colours indicate the area of the worm, see colour scale bars at the bottom of the window.

**Video supplement 5. Segmented worms and their summarized tracks.** The segmented worms are shown in white, whereas the worm outlines at each time point are shown in yellow. The time series shows the individual worms per time point, whereas the yellow outlines are superimposed over the entire time series and shown at each time point of the video.

**Video supplement 6. A segmented worm and its footprint for LR = 11.5.** The orange objects shows the segmented worm at each time point per movie frame, whereas the red area shows the worm’s footprint calculated for the entire time series.

**Video supplement 7. A segmented worm and its footprint for LR = 4.0.** The green object corresponds to the segmented worm shown at each time point, the orange area indicates the footprint of this worm, calculated for the entire time series.

---

**Figure supplement 1.**
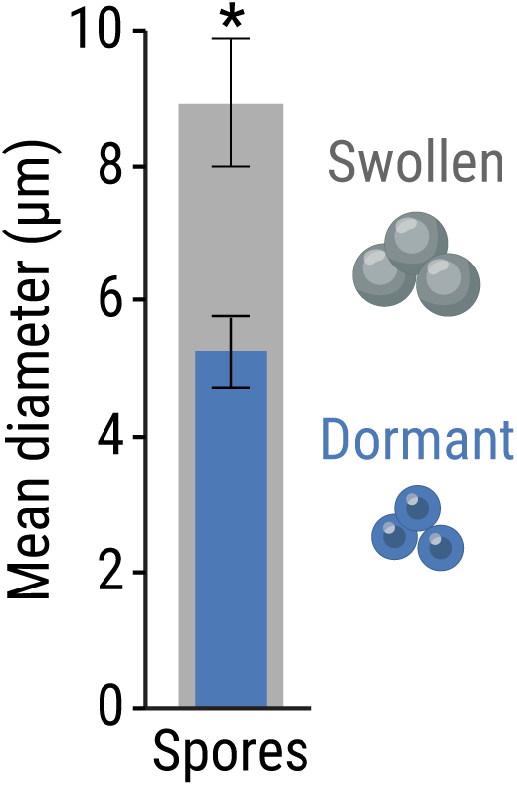
Mean diameter of swollen and dormant *R. microsporus* spores ± one SEM (n = 59 for dormant spores and n = 52 for swollen spores). Unpaired *t*-test with Welch’s correction (\**p<0.05*, Table supplement 3).

**Figure supplement 2.**
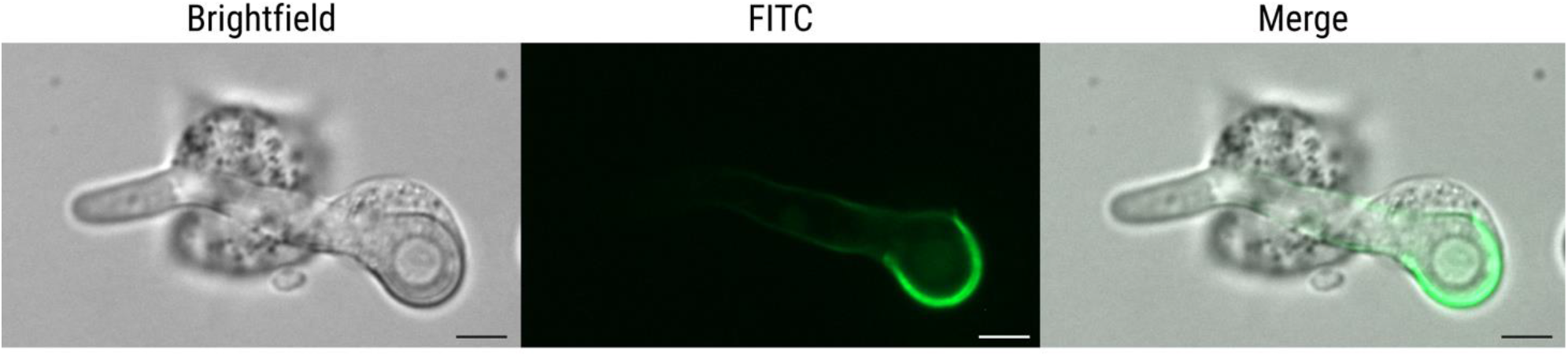
Predation of *Protostelium aurantium* on germinated spores from *Rhizopus microsporus*. Fluorescence microscopy images showing ingestion of a germinating, swollen *R. microsporus* spore (stained with FITC) by *P. aurantium*. Scale bars: 5 μm.

**Figure supplement 3.**
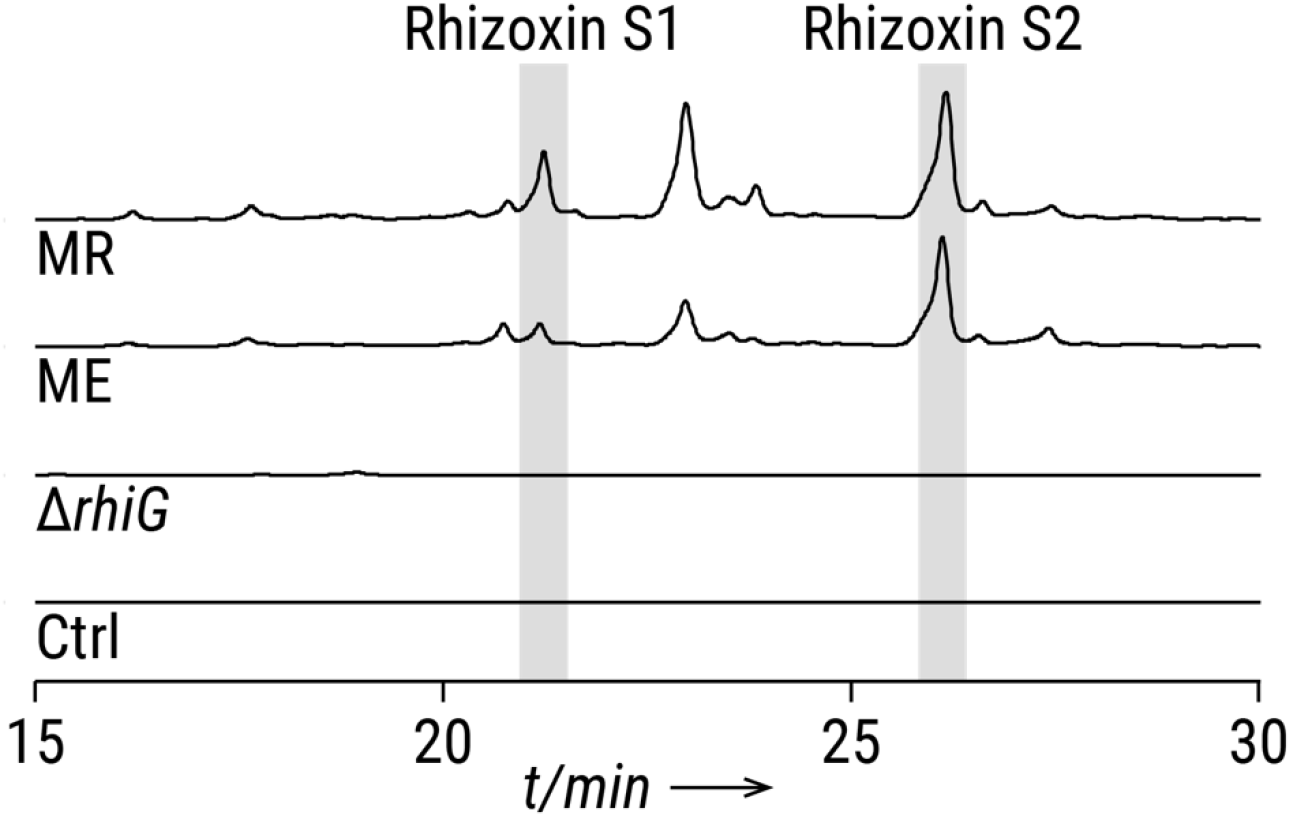
HPLC profiles of crude extracts from axenic endosymbiotic *Mycetohabitans* showing the two major bacterial rhizoxin congeners (rhizoxin S1 and rhizoxin S2). Monitored at 310 nm. MR, *Mycetohabitans rhizoxinica* HKI-0454; ME, *Mycetohabitans endofungorum* HKI-0456; *ΔrhiG*, rhizoxin-deficient mutant; Ctrl, medium control. Cultures were grown to a similar OD_600_ (approx. 3.5). The peak areas were calculated for rhizoxin S1 (MR: 9.07E6, ME: 5.01E6) and rhizoxin S2 (MR: 2.59E7, ME: 2.05E7).

**Figure supplement 4.**
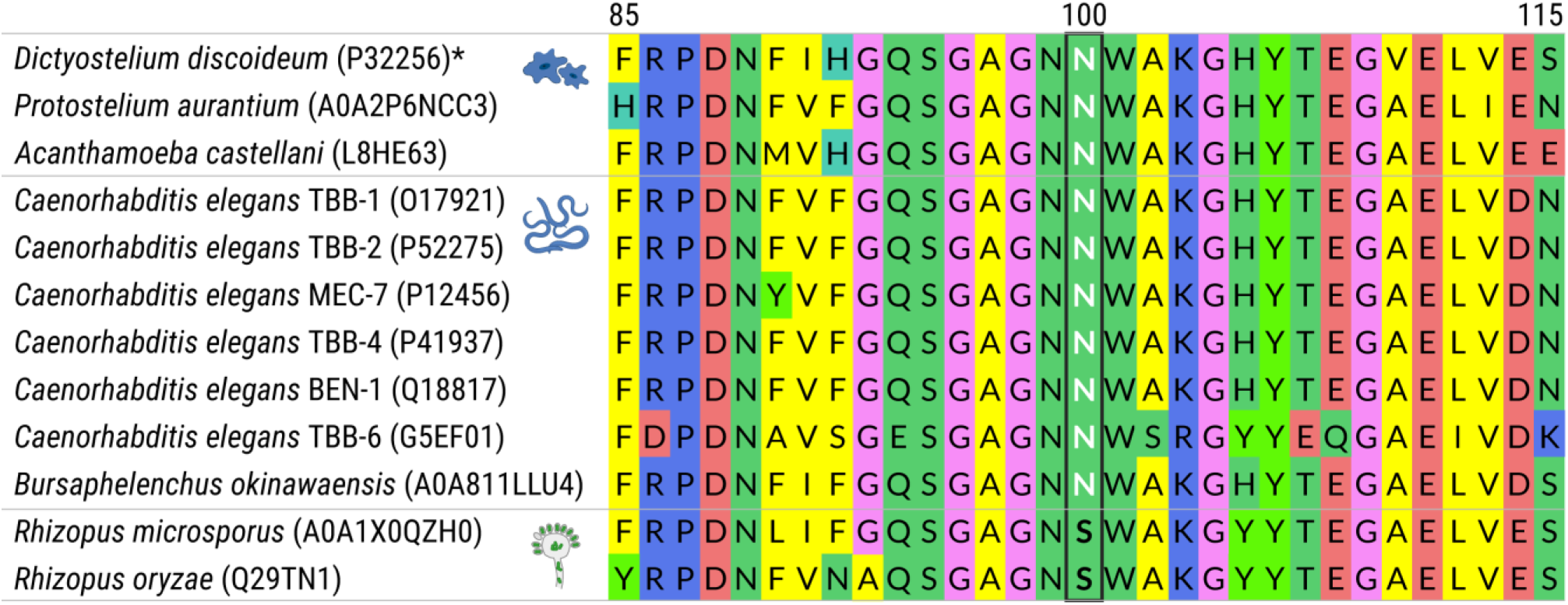
Amino acid sequence alignment of *β-tubulin* proteins from amoebae, nematodes, and *Rhizopus* sp. Numbers refer to the amino acid positions from *Rhizopus microsporus*. A black box at amino acid position 100 indicates the positions of the serine or asparagine that confers rhizoxin resistance (black) or sensitivity (white), respectivel. Accession numbers for proteins retrieved from the Universal Protein Resource are given in brackets. * Indicates protein sequences retrieved from GenBank.

**Figure supplement 5.**
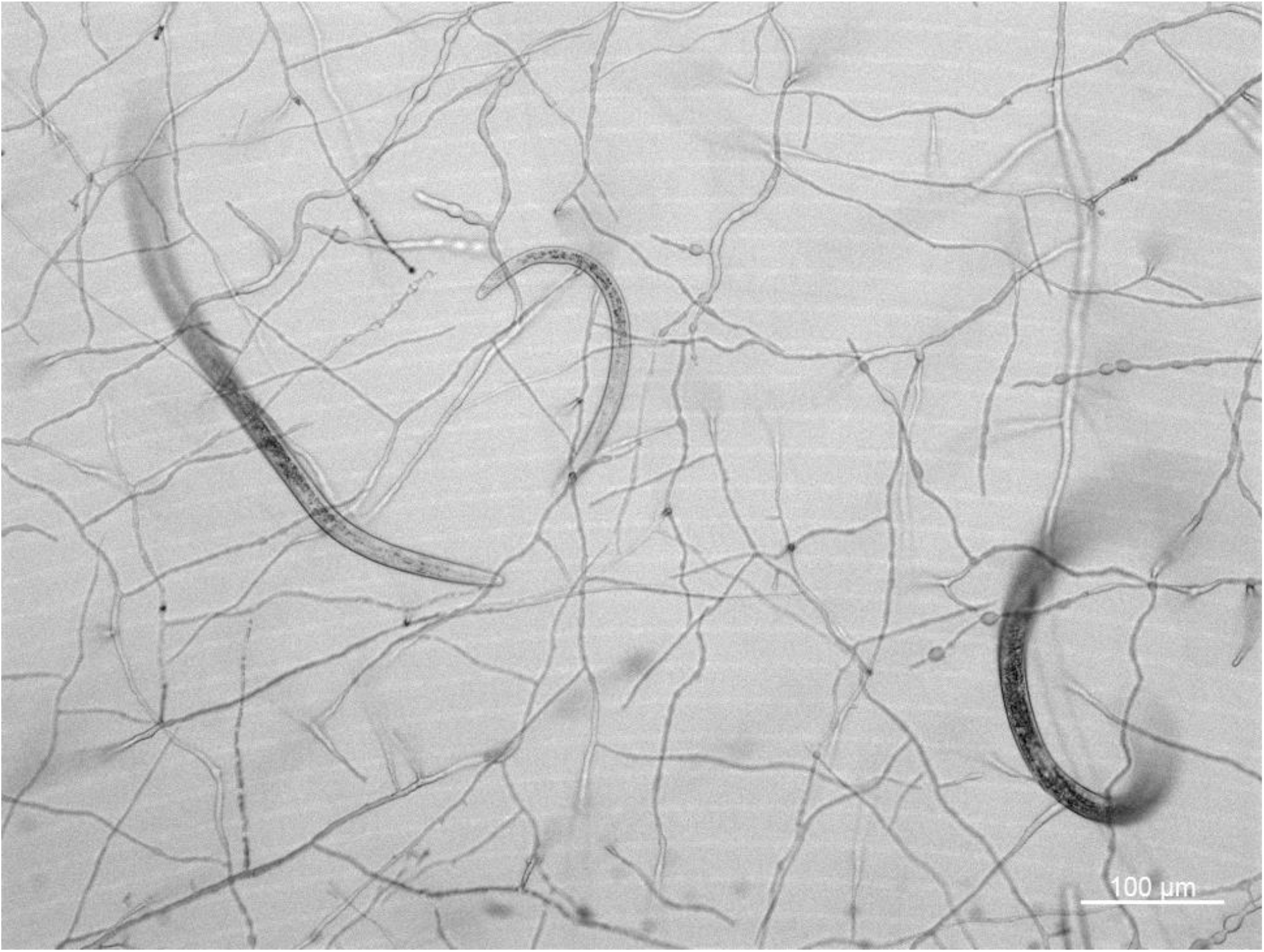
Microscopic image of *Aphelenchus avenae* co-incubated with symbiotic *Rhizopus microsporus*. The majority of worms are dead (Video supplement 2).

**Figure supplement 6.**
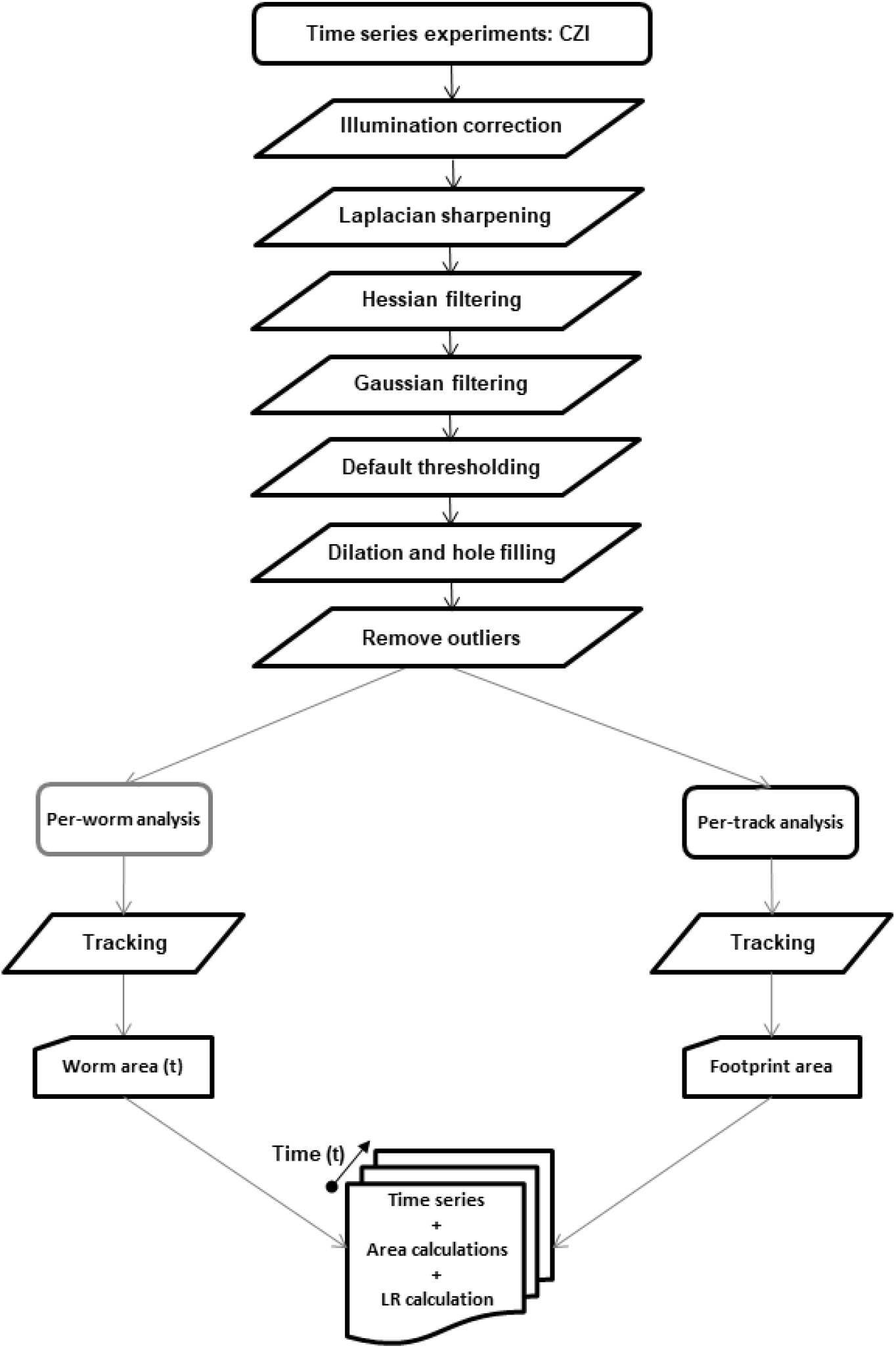
The automated workflow of worm liveliness analysis. The pre-processing and per-timeframe segmentation of images were carried out by using a custom-written JIPipe project (www.jipipe.org). The processing steps were as follows: illumination correction (with 20 pixel Gaussian filtering; Laplacian sharpening (3×3 pixels); Hessian filtering (5 pixels, largest eigenvalue); Gaussian filtering (3 pixels); thresholding (Default method as provided by ImageJ); dilation (1 pixel); morphological hole filling; Remove Outliers (50 pixels size threshold). The resulting images were tracked with JIPipe’s “Split into connected components” algorithm, producing both the per-worm segmented areas and the per-track combined area covered by each worm (the so-called worm footprint). The per-track worm footprints were divided by the per-worm areas at each timepoint, thus providing the so-called Liveliness Ratio (LR).

**Figure supplement 7.**
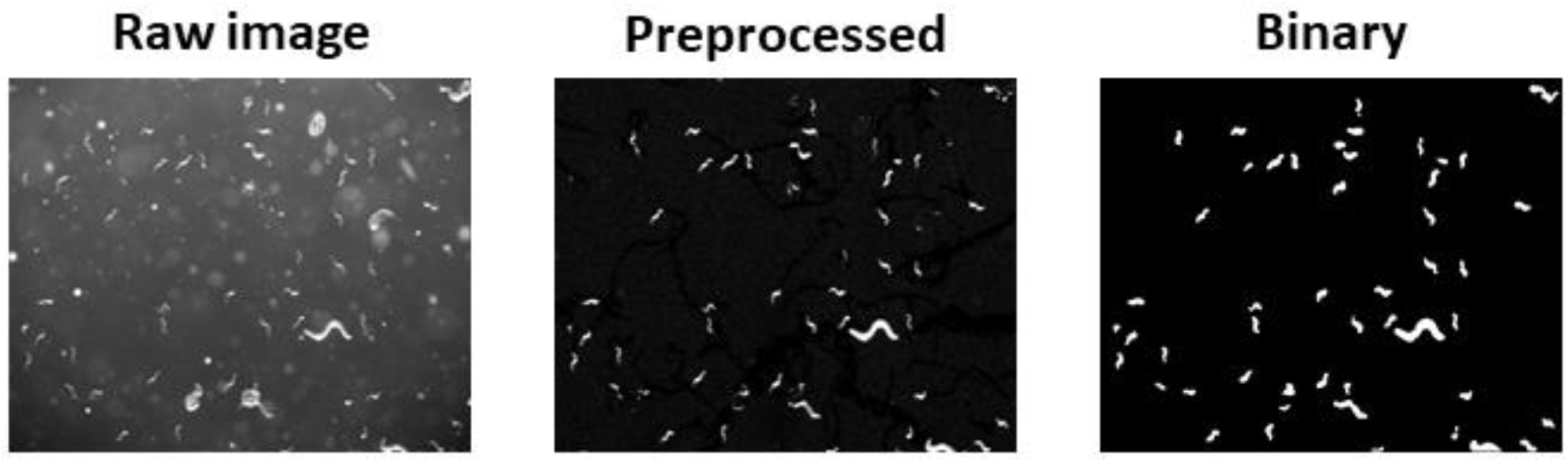
Illustrative steps of the image segmentation process of transmitted light images of live worms. The images were processed according to the workflow described in details in the Methods “Image analysis and mathematical quantification of *Aphelenchus avenae* migration” and in Figure supplement 6. The illustrated steps are: the transmitted light raw image (left); the preprocessed image (after Gaussian filtering); and the binarised image (after Remove Outliers).

**Figure supplement 8.**
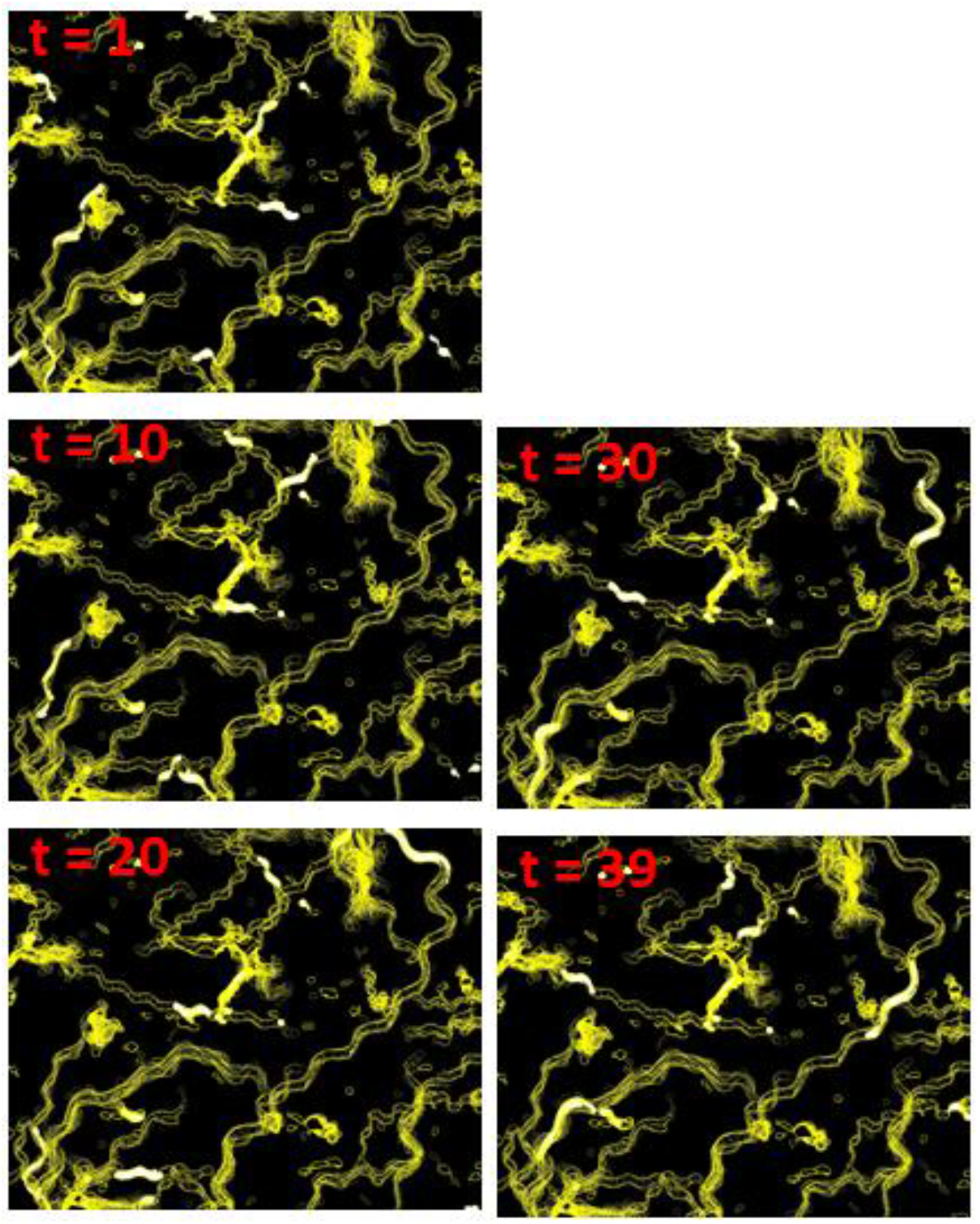
Images from a 39-frame movie showing the summarised outlines of all tracked worms. The yellow lines are the outlines of each tracked worm at every time point. The white elongated objects indicate the segmented worms. The segmentation and tracking were carried out as described in Figure supplement 6. Out of the 39-frame time series of images, here are shown the images at time (t) point 1, 10, 20, 30, and 39. The entire time series of the tracks and segmented worms can be viewed as animation in Video supplement 5.

